# Non-nutritive sweetener consumption during pregnancy affects adiposity in mouse and human offspring

**DOI:** 10.1101/713974

**Authors:** Meghan B. Azad, Alyssa Archibald, Mateusz M. Tomczyk, Alanna Head, Kyle G. Cheung, Russell J. de Souza, Allan B. Becker, Piushkumar J. Mandhane, Stuart E. Turvey, Theo J. Moraes, Malcolm R. Sears, Padmaja Subbarao, Vernon W. Dolinsky

## Abstract

Overweight and obesity affect over 20% of children worldwide. Emerging evidence shows that nonnutritive sweeteners (NNS) could adversely influence weight gain and metabolic health, particularly during critical periods of development. Thus, we aimed to investigate the impact of prenatal NNS exposure on postnatal growth and adiposity. Among 2298 families participating in the CHILD cohort study, children born to mothers who regularly consumed NNS during pregnancy had elevated body mass index and adiposity at 3 years of age. In a complementary study designed to eliminate confounding by human lifestyle factors and investigate causal mechanisms, we exposed pregnant mice and cultured adipocytes to NNS (aspartame or sucralose) at doses relevant to human consumption. In mice, maternal NNS exposure caused elevated body weight, adiposity and insulin resistance in offspring, especially in males. Further, in 3T3-L1 pre-adipocyte cells, sucralose exposure during early stages of differentiation caused increased lipid accumulation and expression of adipocyte differentiation genes (e.g. C/EBP-α, FABP4, FAS). The same genes were upregulated in the adipose tissue of male mouse offspring born to sucralose-fed dams. Together, these clinical and experimental findings provide evidence suggesting that maternal NNS consumption induces obesity risk in the offspring through effects on adiposity and adipocyte differentiation.

**One Sentence Summary:** Maternal consumption of non-nutritive sweeteners during pregnancy stimulates adipocyte differentiation, insulin resistance, weight gain, and adiposity in mouse and human offspring.

## Introduction

Globally, over 20% of children are overweight or obese, with rates exceeding 50% in some countries (1). The obesity epidemic has arisen alongside a surge in over-nutrition, which stimulates adipocytes to expand and store excess calories. Mounting evidence shows that obesity originates early in life, perhaps even *in utero*. The Developmental Origins of Health and Disease (DOHaD) hypothesis postulates that prenatal and early postnatal exposures can “program” lifelong metabolism, weight gain, and other endocrine pathways (reviewed in (2)). Moreover, in early life, environmental exposures can stimulate adipocyte precursor cells to induce the process of adipocyte differentiation, creating a large reservoir of adipocytes to support the development of obesity in response to over-nutrition later in life (3-7).

Excess energy intake from sugar, especially sugar-sweetened beverages, is strongly associated with obesity (8-11); hence, sugar substitutes or “non-nutritive sweeteners” (NNS) including aspartame and sucralose are marketed as healthier alternatives (12, 13). NNS are widely consumed, including by pregnant women. Almost 30% of mothers in the Canadian CHILD cohort consumed NNS during pregnancy (14), and similar rates have been reported in the USA (24% (15)) and Denmark (45% (16)). Contrary to their intended benefits, NNS have been inconsistently associated with metabolic derangements and adverse effects on cardiometabolic health in adults (17-19) and children (20); however, few studies have investigated the metabolic effects of NNS exposure *in utero*. In the CHILD cohort, we found that daily NNS beverage consumption during pregnancy was associated with a 2-fold higher risk of infant overweight at 1 year of age, compared to no maternal NNS beverage consumption (14). Similar results were observed among older children in the Danish National Birth Cohort (16). Interestingly, both studies observed stronger effects in males, although a third study found no association in US children of either sex (21).

Limited evidence from animal studies also suggests that NNS consumption during pregnancy and lactation may predispose offspring to develop obesity and metabolic syndrome (22). However, most studies have used doses that exceed the human acceptable daily intake (ADI), which is equivalent to at least 20 packets of NNS or 12 cans of diet soda per day for the average person (13). In mice, Collison et al. found that chronic lifetime exposure to NNS (55 mg/kg/day aspartame; exceeding the ADI by 1.4-fold), commencing *in utero*, was associated with increased weight gain and decreased insulin sensitivity in adulthood (23), but the impact of maternal NNS intake was unclear because exposure was maintained in the offspring after weaning. A more recent study by Olivier-Van Stichelen et al. found that maternal NNS intake (a combination of sucralose and acesulfame-K at 2-fold ADI) altered the microbiome and metabolism of young offspring and *reduced* their total body weight (24), although adiposity was not assessed and the offspring were not followed beyond the weaning period. Von Poster Toigo et al. found that rats exposed to an even higher dose of NNS (343 mg/kg/day aspartame; 8.6-fold ADI) during gestation gain more weight and have altered lipid profiles during adulthood (25), yet other studies have reported no difference in weight gain following prenatal NNS exposure (26). Similarly, conflicting evidence from *in vitro* studies suggests that NNS can either stimulate (27) or downregulate (28, 29) adipocyte differentiation.

Overall there is a paucity of evidence from human and experimental studies on the potential impact of prenatal NNS exposure on the development of obesity and metabolic health. Here, we extend our previous findings on maternal NNS beverage consumption and infant body composition in the CHILD cohort (14) by re-assessing this relationship at 3 years of age. Further, we use experimental model systems to examine the underlying mechanisms in mice, using physiologically relevant doses of both aspartame and sucralose. Finally, we characterized these mechanisms using an *in vitro* model of adipocyte differentiation and show for the first time that sucralose exerts effects on adiposity and adipocyte differentiation. The combination of clinical and experimental findings provides new evidence that maternal NNS consumption conditions obesity risk in the offspring.

## Results

### Maternal NNS intake during pregnancy is associated with higher BMI and adiposity in children

Building on our previous findings at 1 year of age in the CHILD pregnancy cohort (14), we re-examined the association of maternal NNS consumption and child body composition at 3 years of age among 2298 mother-child dyads. During pregnancy, 29.9% of mothers reported consuming any NNS beverages and 5.2% consumed them daily (**Table S1**). Consistent with our previous results, children born to mothers reporting daily NNS beverage consumption had significantly higher BMIs at 3 years of age than children born to mothers who did not consume NNS beverages (mean z-score 0.88 ± 0.97 vs. 0.53 ± 0.96; crude β = 0.37, 95% CI 0.19 – 0.55), although this association was partially attenuated after adjusting for potential confounders including maternal BMI, diabetes, smoking and overall diet quality, child diet quality and screen time (aβ = 0.17, 95% CI −0.05 – 0.39) (**Fig. 1** and **Table S2**). Similar results were observed for girls and boys, and for the adiposity outcome of subscapular skin folds (**Fig. 1** and **Table S3**). Together, these results suggest that maternal NNS consumption during pregnancy may promote excessive weight gain or adiposity in offspring, although confounding by lifestyle factors appears to partially explain this relationship. To eliminate the possibility of confounding, determine causality and investigate biological mechanisms, we undertook mechanistic studies exposing pregnant mice and cultured adipoctyes to NNS at doses relevant to human consumption.

**Figure 1.**
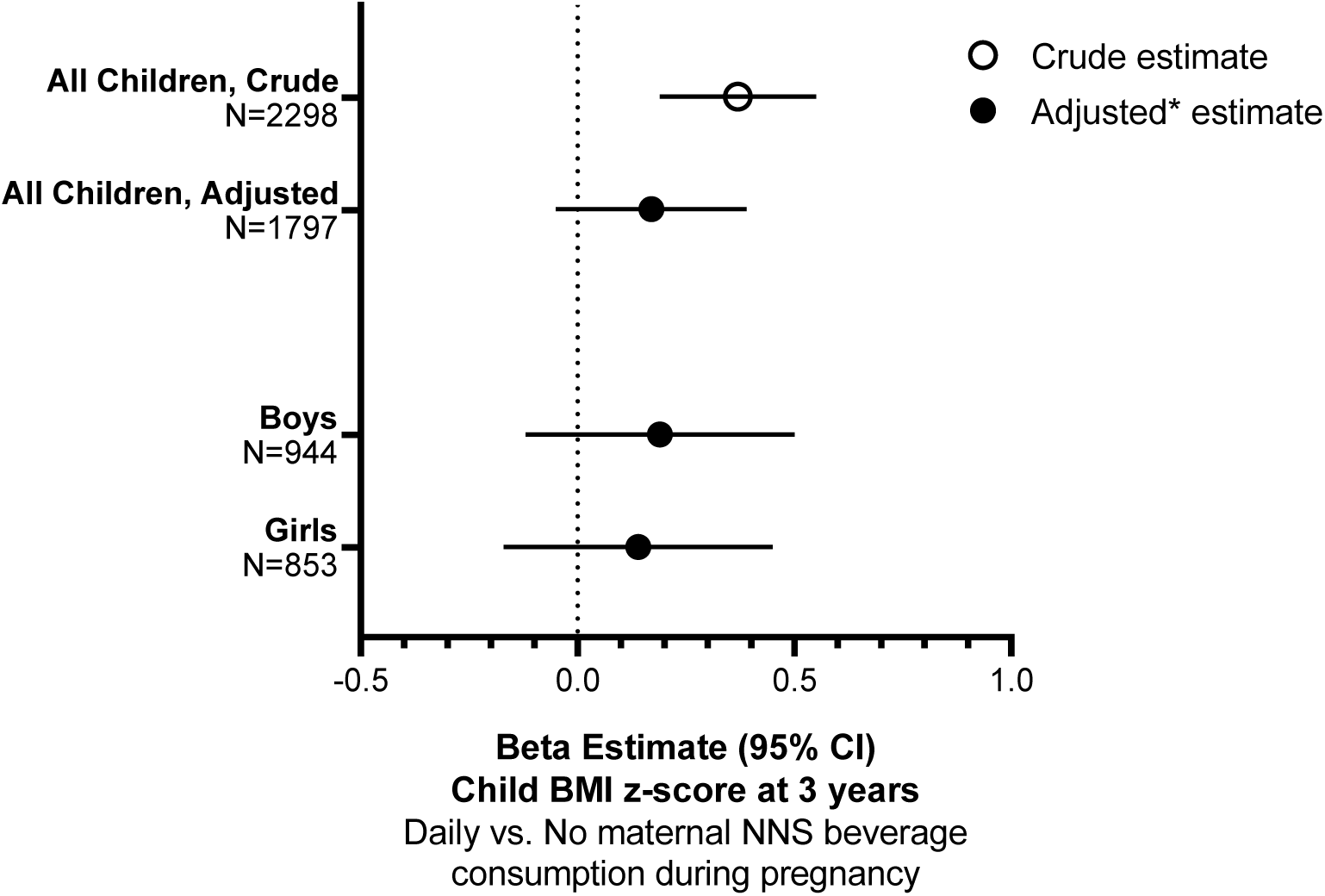
Maternal consumption of NNS-sweetened beverages and child body mass index (BMI) at 3 years of age in the CHILD cohort. Estimates for highest consumption group (≥ 1 beverage per day) vs no consumption. *Adjusted for maternal BMI, total energy intake, Healthy Eating Index score, sugar sweetened beverage intake, postsecondary education, smoking and diabetes during pregnancy; breastfeeding duration; child sex, screen time, fresh and frozen food intake, and soda consumption. CI, confidence intervals; NNS, non-nutritive sweetener.

### Sucrose and NNS variably impact weight gain and energy intake in pregnant mice

At e18.5, pregnant mice receiving sucrose in their drinking water weighted more than the control dams, although this difference did not reach statistical significance (**Table 1**). Dams receiving sucrose consumed more sweetened water and more food, thus their average daily energy intake was ∼1.4-fold greater than controls (**Table 1**; p<0.01). Dams receiving aspartame or sucralose increased their food intake by a lesser degree (1.1 and 1.2 fold, respectively; p<0.01), but their body weight was not affected. Notably, maternal sucrose and aspartame consumption increased the number of pups in the litters (**Table 1**; p<0.01) whereas sucralose did not significantly affect the litter size.

**Table 1.** Maternal mouse parameters.

|  | Control<br>(Plain Water) | Sucrose<br>(45 g/L) | Aspartame<br>(0.2 g/L) | Sucralose<br>(0.04 g/L) |
| --- | --- | --- | --- | --- |
| Body weight at e18.5 (g) | 33.00 ± 0.38 | 34.80 ± 0.48 | 30.23 ± 1.15 | 32.33 ± 1.14 |
| Water consumption (ml/day) | 4.32 ± 0.25 | 8.09 ± 1.41* | 6.28 ± 0.43 | 5.08 ± 0.44 |
| Mean NNS dose (mg/kg/day) | N/A | N/A | 41.55 | 6.29 |
| Human ADI for comparison (mg/kg) | N/A | N/A | 40 | 5 |
| Food consumption (g/day) | 5.57 ± 0.06 | 7.70 ± 0.25** | 6.35 ± 0.06** | 6.84 ± 0.12** |
| Energy intake (kcal/day) | 94.7 ± 2.1 | 130.8 ± 15.2* | 107.9 ± 2.2 | 116.3 ± 7.0 |
| Energy intake (kcal/day/kg) | 3591 ± 231 | 4949 ± 686 | 4797 ± 334 | 4600 ± 180 |
| Litter Size | 5.67 ± 0.50 | 8.33 ± 0.19** | 7.33 ± 0.19** | 6.33 ± 0.41 |

n=6 dams per group. Values are mean ± SEM. Comparisons by One-way ANOVA with Bonferroni post-test. \*p<0.05;
\*\*p<0.01 vs. Control.

### Maternal NNS intake has sex-specific effects on adiposity in mouse offspring

Next, we investigated the influence of maternal NNS intake on weight gain in male and female mouse offspring. Maternal sucrose, aspartame and sucralose consumption all conditioned increased body weight in male offspring by 7 weeks of age compared to the male offspring of control dams (**Fig. 2A**; all p<0.001). The elevated body weight in these male offspring persisted until sacrifice at 11 weeks of age. Conversely in female offspring, only maternal sucrose consumption (not aspartame or sucralose) induced elevated body weight at 10 and 11 weeks of age (**Fig. 2B**; p<0.05). Interestingly, the elevated body weight did not appear to be due to differences in energy intake because average daily food intake was similar across all offspring groups **(Table S4)**.

**Figure 2.**
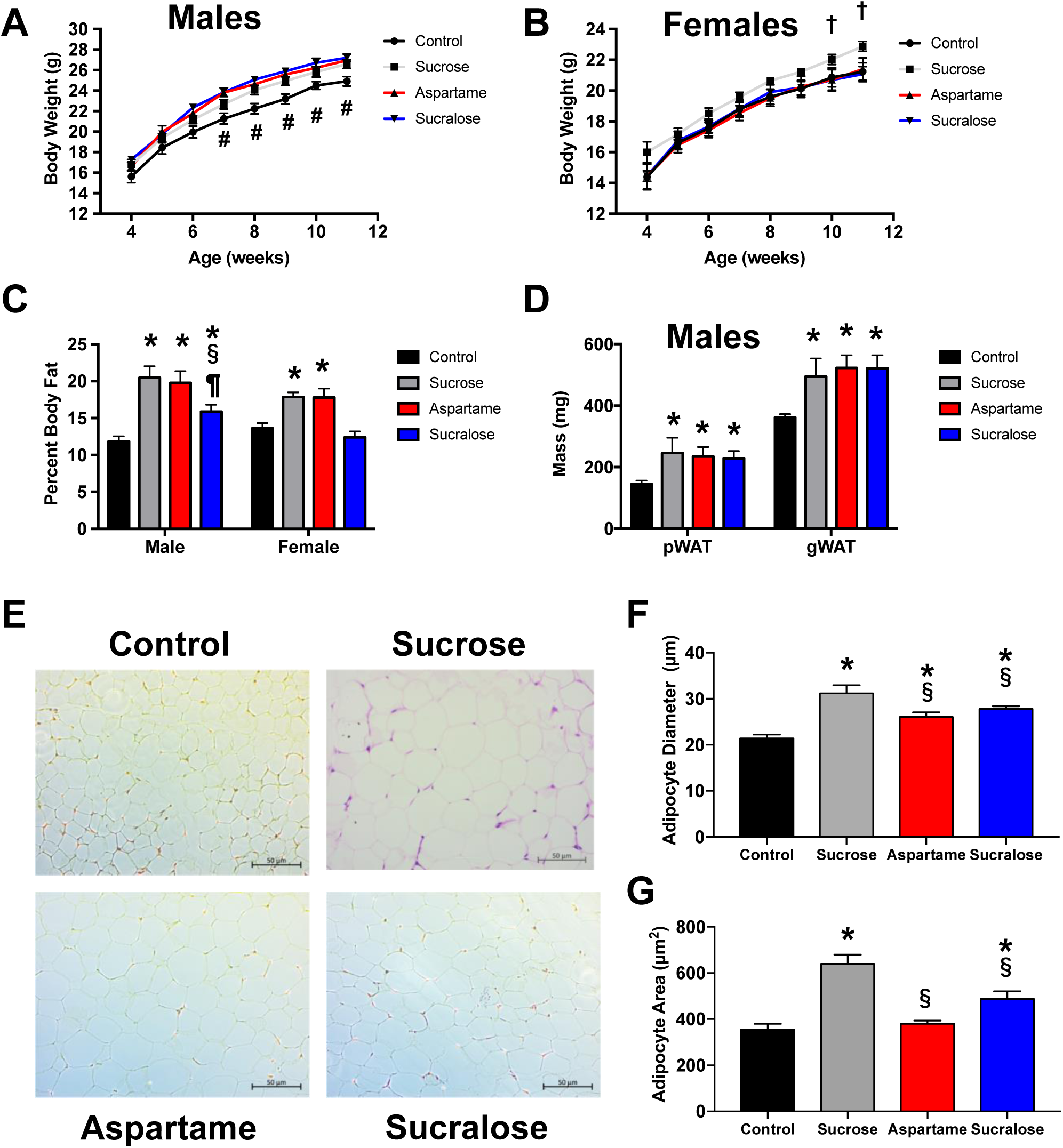
Body composition in male and female offspring of dams fed sucrose, aspartame or sucralose during pregnancy and lactation. Body weight trajectory of (A) male and (B) female offspring; (C) Percent body fat of male and female offspring at 11 weeks of age; (D) pWAT and gWAT weights of male offspring; (E) Representative images of H&E stained sections of pWAT adipocytes at 20x magnification; Adipocyte (F) diameter and (G) area, values represent mean +/− SEM, n=6. #significant differences (p<0.05) between control male offspring vs. the male offspring of sucrose, aspartame and sucralose dams as calculated by a two-way repeated measures ANOVA with a Bonferroni post-test. †significant differences (p<0.05) between female offspring of sucrose dams vs. the offspring of control, aspartame and sucralose dams by a two-way repeated measures ANOVA with a Bonferroni post-test. Significance after one-way ANOVA with Bonferroni post-hoc tests: *p <0.05 vs. offspring of control dams, §p<0.05 vs. offspring of sucrose dams and ¶p<0.05 vs. offspring of aspartame dams. pWAT: Perirenal White Adipose Tissue, gWAT: Gonadal White Adipose Tissue.

To determine whether increased body weight was related to alterations in lean and/or fat mass in the offspring, we performed dual energy X-ray absorptiometry (DXA). This analysis showed that maternal sucrose, aspartame and sucralose all markedly increase the percent body fat (50%, 47%, and 15% increases, respectively) in male offspring, compared to controls (**Fig. 2C**; p<0.0001). Maternal sucrose and aspartame also increased the percent body fat in female offspring (**Fig. 2C**; p<0.05); however, sucralose had no effect on percent body fat in females. Consistent with these observations, maternal consumption of sucrose, aspartame and sucralose all increased the weight of perirenal white adipose tissue (pWAT) and gonadal white adipose tissue (gWAT) fat pads of the male offspring, compared to controls (**Fig. 2D**; p<0.05). Notably, the effect of sucralose was dose dependent, as lower levels of sucralose administration to dams did not induce elevated body weight and fat pad mass in the male offspring **(Table S5)**. H&E staining of perirenal adipose tissue (**Fig. 2E**) revealed that maternal aspartame and sucralose consumption increased the mean adipocyte diameter of the male offspring by 22% and 30%, respectively (**Fig. 2F**, p<0.05). Adipose tissue was the only major organ system that increased in weight; the liver, heart, kidney and spleen of the offspring were generally similar between all groups **(Table S6)**. One notable exception was increased liver mass in female offspring of sucrose-fed dams **(Table S6)**.

### Maternal NNS intake has sex-specific effects on insulin sensitivity in mouse offspring

Next, since maternal NNS consumption increased body fat accumulation in offspring, we examined whether insulin sensitivity was also affected. Glucose tolerance tests of the male offspring did not show any differences across groups (**Fig. 3A, B**). In the female offspring, maternal sucrose consumption induced significant glucose intolerance, while maternal aspartame and sucralose consumption had no effect (**Fig. 3C, D**). Insulin tolerance tests revealed that the male offspring of sucrose, aspartame and sucralose-fed dams were more insulin resistant than the male offspring of control dams (**Fig. 3E, F**). However, the insulin sensitivity of the female offspring was similar across all groups (**Fig. 3G, H**).

**Figure 3.**
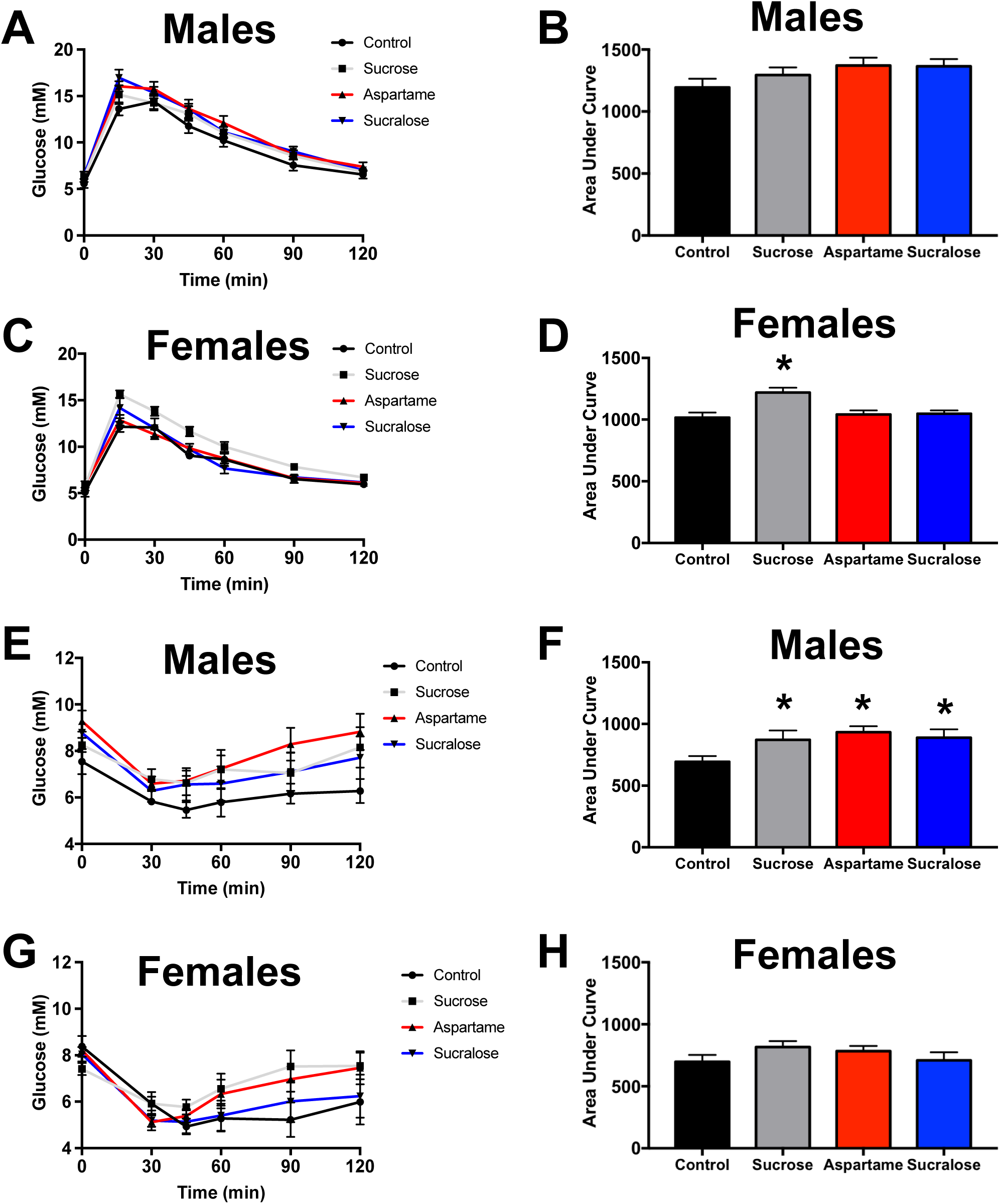
Glucose tolerance and insulin sensitivity in 10-week old male and female offspring of dams fed sucrose, aspartame or sucralose during pregnancy and lactation. (A) GTT and (B) area under the curve in male offspring; (C) GTT and (D) area under the curve in female offspring; (E) ITT and (F) area under the curve in male offspring; (G) ITT and (H) area under the curve in female offspring. GTT: Glucose Tolerance Test, ITT: Insulin Tolerance Test. Values represent the mean +/− SEM, n=6. *p-values represent significant differences (p<0.05) vs. control offspring as calculated by one-way ANOVA with Bonferroni post-hoc tests.

### Sucralose has pro-adipogenic effects on 3T3-L1 pre-adipocytes in vitro

Since maternal NNS influenced body fat accumulation in male mouse offspring, and early-life is a critical stage that determines stem cell fate, we examined the effects of sucralose in cultured cells using the well-established male 3T3-L1 pre-adipocyte cell line. Previous research has shown that aspartame affects lipid accumulation and adipocyte differentiation in 3T3-L1 cells (29). Therefore, we examined the stage(s) of adipocyte differentiation affected by sucralose. To this end, the 3T3-L1 adipocytes were incubated with induction medium in the presence or absence of sucralose (200nM) for the indicated periods of time, as illustrated in **Fig. 4A**. As expected, control cells incubated with induction medium for 8 days differentiated into adipocytes, exhibited by lipid accumulation as judged by oil red staining (**Fig. 4B**, treatment a). Cells treated with sucralose from d0-d2 (treatment b, modeling germline exposure) or d0-d8 (treatment e, throughout differentiation) exhibited the highest accumulation of lipid (**Fig. 4B**). Of note, lipid accumulation was not significantly affected by sucralose treatment in other time windows, including d2-d4 (treatment c, modeling fetal exposure) or d4-d8 (treatment d, modeling postnatal exposure) **Fig. 4B**). These results collectively suggest that sucralose administration enhances adipogenesis at an early phase of differentiation, consistent with the effects of prenatal NNS exposure on body fat accumulation observed in mice (**Fig. 2**) and 3 year-old participants in the CHILD study (**Fig. 1**).

**Figure 4.**
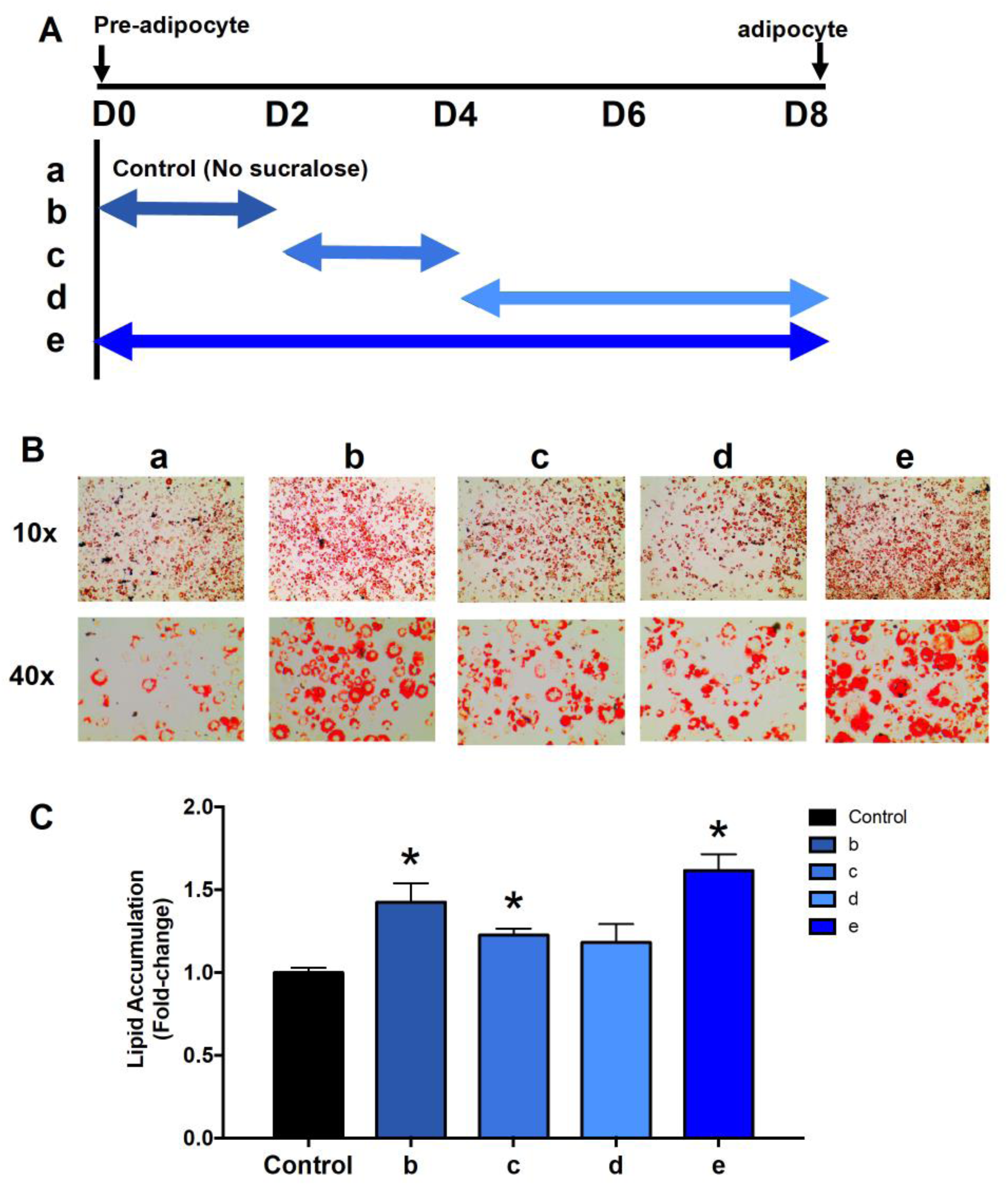
Effect of sucralose on 3T3-L1 adipocyte differentiation. (A) Schematic outline of the experimental design: double-headed arrows indicate the length of treatment. 3T3-L1 cells were treated with 200nM sucralose for the indicated periods of time. (B) Oil red O staining to measure cellular lipid content assessed 8 days following induction of 3T3-L1 pre-adipocyte differentiation with induction medium containing MDI, insulin and fetal bovine serum in the presence or absence of 200nM sucralose. (C) Quantification of cellular lipid content. Values represent the mean +/− SEM of data from 3 independent experiments with 3 replicates. *p<0.05 vs. control (no sucralose treatment) as calculated by one-way ANOVA with Bonferroni post-hoc tests.

### Sucralose stimulates pro-adipogenic regulators and enzymes in vitro and in vivo

Since adipocyte differentiation is a complex process that can be modulated by multiple stimuli including transcription factors, we examined how sucralose affected the gene expression of regulators of the adipocyte phenotype. The addition of sucralose to the culture media from d0-d8 (treatment e) induced a small but significant increase in the expression of the peroxisomal proliferator activated receptor (PPAR)-γ transcription factor at d8 of adipocyte differentiation (*Pparg*; **Fig. 5A**). On the other hand, the addition of sucralose earlier in the adipocyte differentiation program as well as throughout (treatments c and e), induced marked increases in the expression of the adipogenesis-dependent transcription factor, CCAT enhancer binding protein (C/EBP)-α by d8 of adipocyte differentiation (*Cebpa*; **Fig. 5B**). Moreover, the addition of sucralose to the media at the early stages of differentiation (treatments b and c) as well as throughout (treatment e), induced 1.5 to 2-fold increases in the mRNA expression of the adipocyte marker genes, adiponectin (*Adipoq*; **Fig. 5C**) and fatty acid binding protein (*Fabp4*; **Fig. 5D**). Consistent with these findings, sucralose also increased the expression of the lipid droplet coat protein, perilipin (*Plin2*; **Fig. 5E**). Sucralose did not affect the expression of the adipogenesis inhibitory factor, *Pref-1* (**Fig. 5F**), suggesting that most of the effects of sucralose are driven by promoting adipogenesis rather than removing factors that maintain the undifferentiated state. Overall, treatment of the cells with sucralose at earlier stages of adipocyte differentiation had remarkable effects on regulators of the adipocyte phenotype whereas treatment of the cells with sucralose at later stages had no effect.

**Figure 5.**
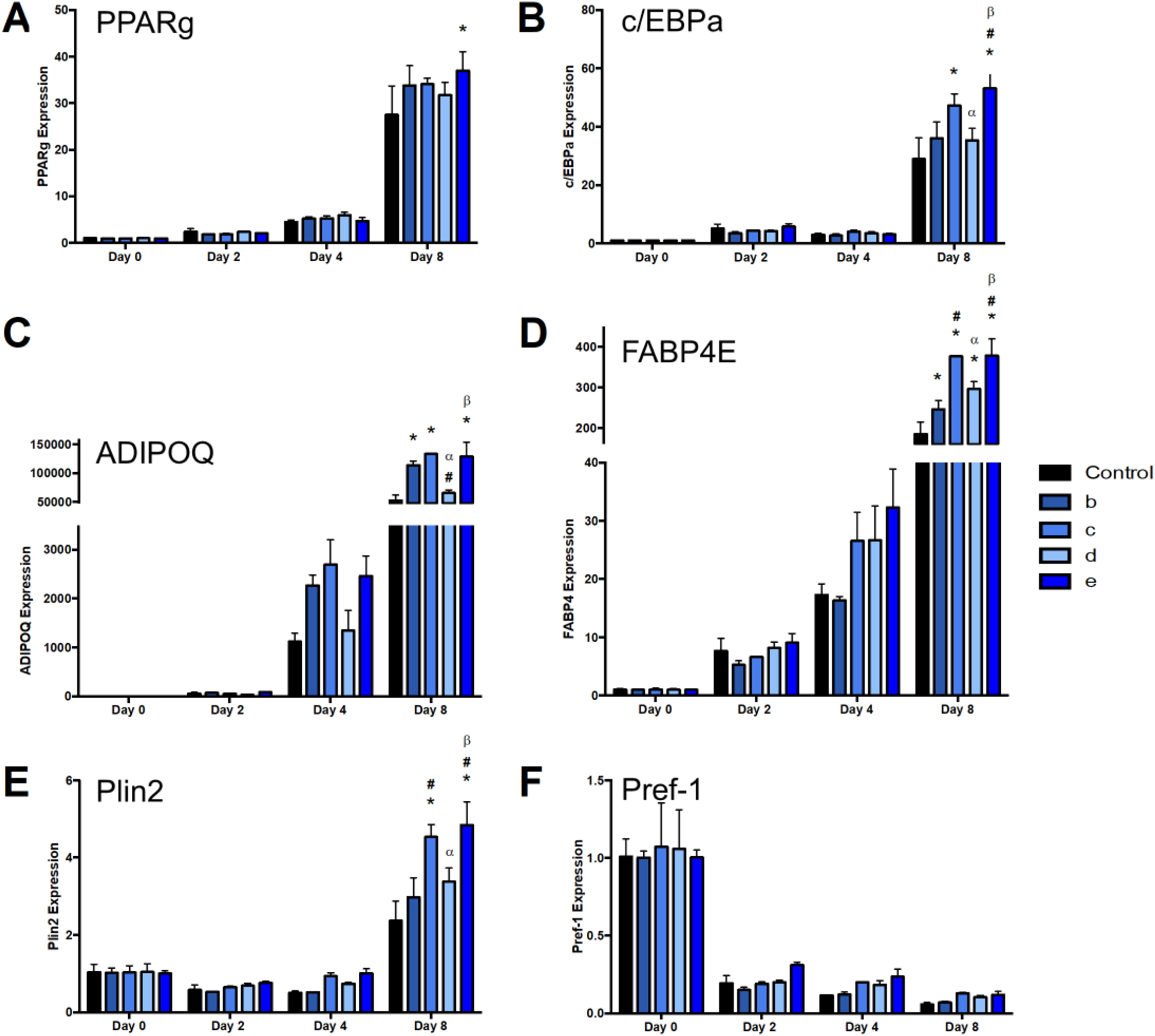
Sucralose increases the expression of pro-adipogenic regulators in 3T3-L1 cells. (A) *pparg* gene expression, (B) *Cebpa* gene expression, (C) *Adipoq* gene expression. (D) *Fabp4* gene expression, (E) *Plin1* gene expression, (F) *Pref1* gene expression. Values represent the mean +/− SEM of data from 3 independent experiments with 3 replicates. qPCR gene expression is relative to the geomean of *Eif2a* and *CycA* and normalized the control group. *p<0.05 vs. control (no sucralose treatment), ^#^p<0.05 vs. b, ^α^p<0.05 vs. c, ^β^p<0.05 vs. d, as calculated by two-way ANOVA with Bonferroni post-hoc tests.

Next, we examined whether sucralose also impacted the expression of genes encoding metabolic enzymes involved in fat storage and mobilization during adipocyte differentiation. Indeed, sucralose administration early in the adipocyte differentiation program increased the expression of fatty acid synthase (*Fasn*; **Fig. 6A**) as well as glycerol phosphate acyltransferase (*Gpam*; **Fig. 6B**). Sucralose also significantly increased the expression of hormone sensitive lipase (*Lipe*; **Fig. 6C**) and adipose tissue triglyceride lipase (*Atgl*; **Fig. 6D**). These findings suggest that sucralose promotes fatty acid and triacylglycerol synthesis as well as its mobilization in differentiating 3T3-L1 adipocytes.

**Figure 6.**
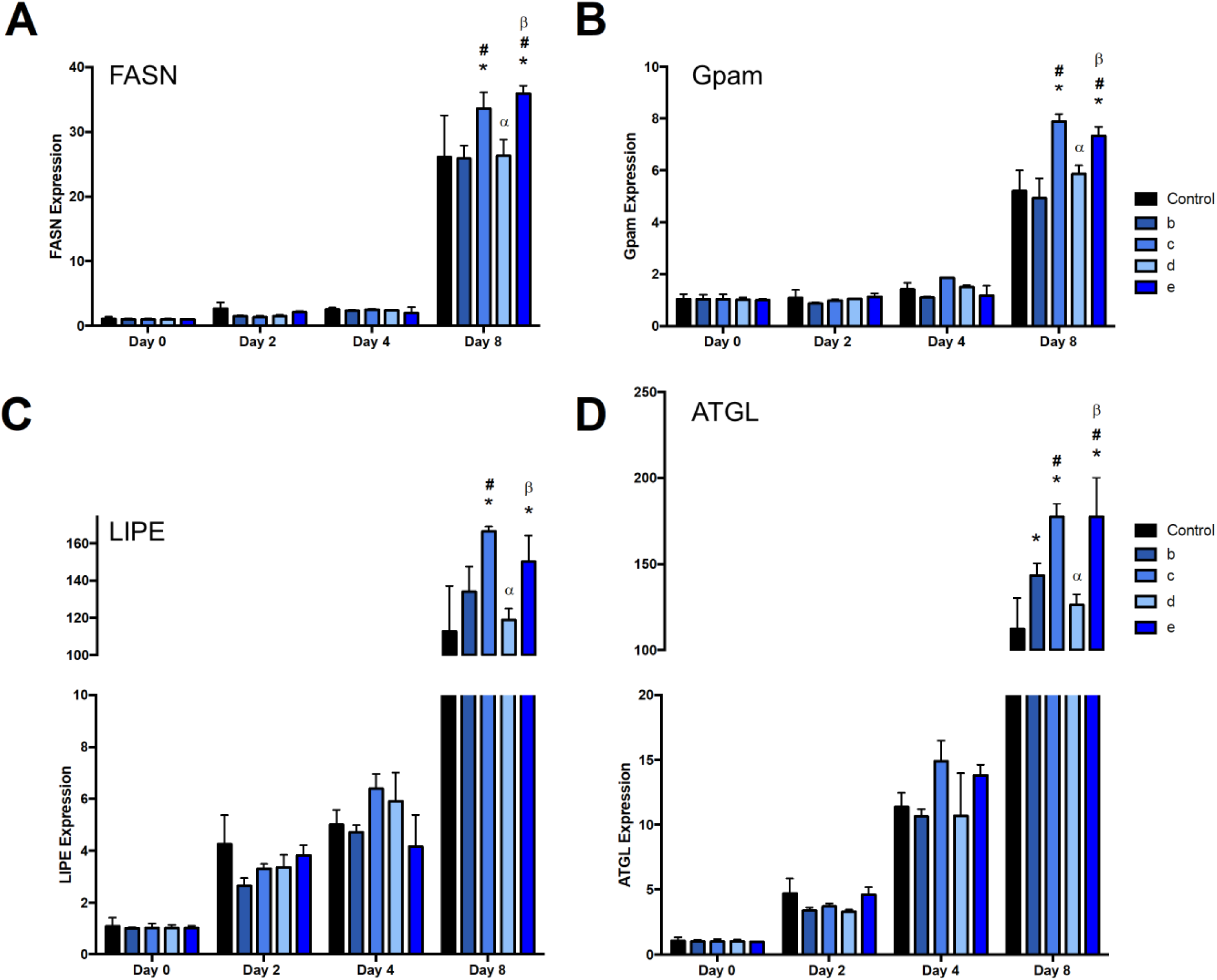
Sucralose stimulates expression of fat storage and mobilization genes in 3T3-L1 cells. (A) *Fasn* gene expression, (B) *Gpam* gene expression, (C) *Lipe* gene expression. (D) *Atgl* gene expression. Values represent the mean +/− SEM of data from 3 independent experiments with 3 replicates. qPCR gene expression is relative to the geomean of *Eif2a* and *CycA* and normalized the control group. *p<0.05 vs. control (no sucralose treatment), ^#^p<0.05 vs. b, ^α^p<0.05 vs. c, ^β^p<0.05 vs. d, as calculated by two-way ANOVA with Bonferroni post-hoc tests.

Finally, we assessed whether maternal sucralose consumption also affected the expression of several of these genes in the pWAT of male mouse offspring. Interestingly, in the offspring of sucralose-fed dams, as well as sucrose-fed and aspartame-fed dams, a ∼1.5-fold increase *Cebpa* and *Fabp4* mRNA expression in pWAT was observed compared to the offspring of control dams (**Fig. 7A, B**). In addition, sucralose (but not sucrose or astpartame) increased *Fasn* and *Gpam* mRNA expression ∼7-fold and ∼3-fold, respectively, compared to the offspring of control dams (**Fig. 7C, D**).

**Figure 7.**
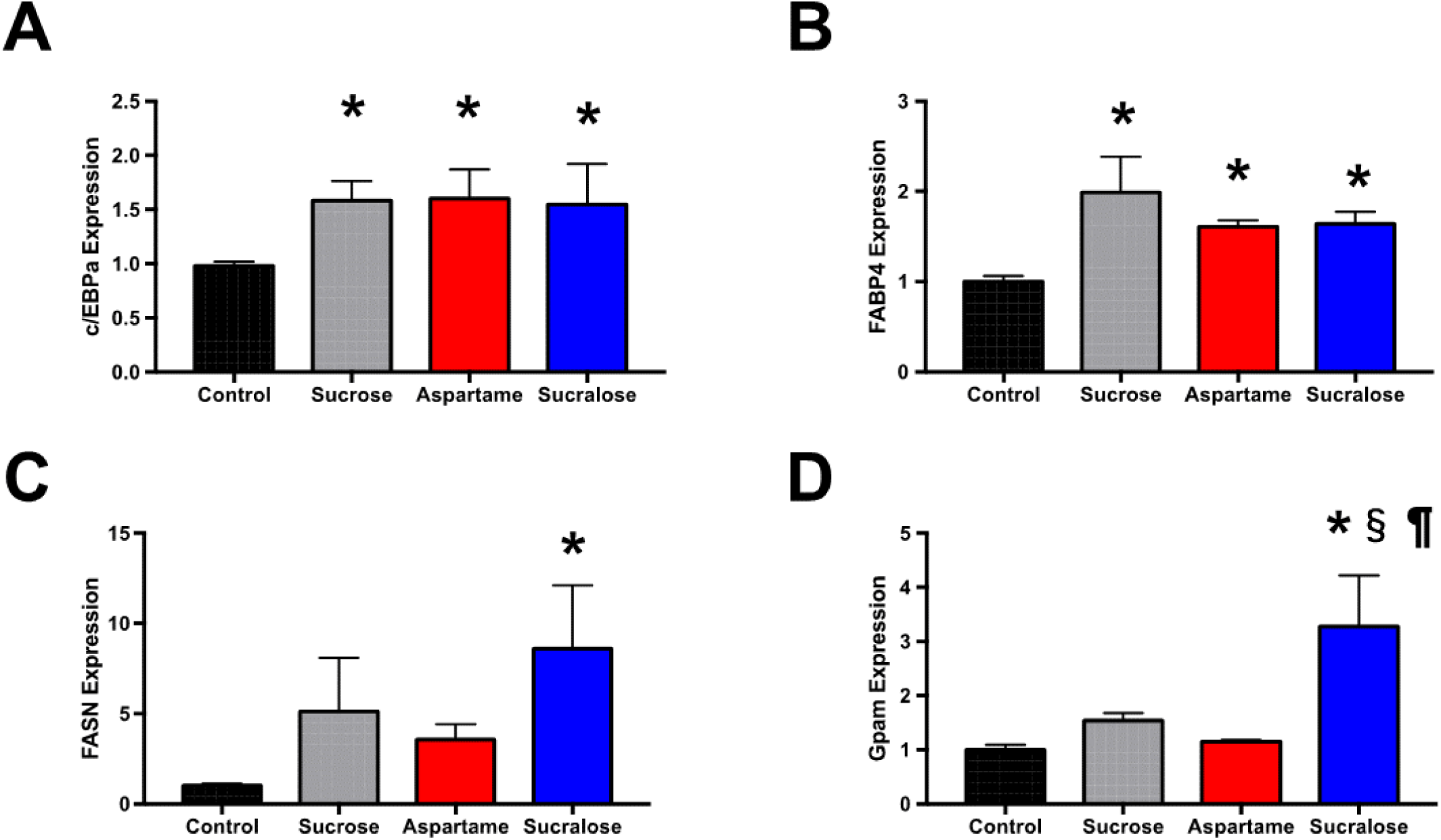
Sucralose increases the expression of pro-adipogenic regulators and fat storage and mobilization genes in murine offspring adipose tissue *in vivo*. (A) *Fasn* gene expression, (B) *Gpam* gene expression, (C) *Lipe* gene expression. (D) *Atgl* gene expression. Values represent the mean +/− SEM, n=6. qPCR gene expression is relative to the geomean of *Eif2a* and *CycA* and normalized the control group. p-values that represent significance after one-way ANOVA with Bonferroni post-hoc tests: *p <0.05 vs. control offspring dams, §p<0.05 vs. offspring of sucrose dams and ¶p<0.05 vs. offspring of aspartame dams.

## Discussion

Our study provides new evidence on the potential adverse effects of NNS, which are typically marketed as ‘healthier’ alternatives to caloric sweeteners, especially for the purposes of weight management and diabetes control. Given that maternal obesity and gestational diabetes are on the rise (30), NNS may be especially appealing to pregnant women, yet very few studies have explored the long-term impact of NNS exposure *in utero*. Here, we used a translational approach to triangulate evidence from a human cohort, a mouse model, and cell culture experiments to show that prenatal NNS exposure influences adipocyte differentiation, fat mass accumulation, and adiposity in offspring.

In the prospective CHILD cohort, we found that children born to mothers who regularly consumed NNS-sweetened beverages had higher BMI and adiposity by three years of age. This association was attenuated after adjusting for maternal BMI and other confounders, which cannot be fully disentangled in an observational study. Thus, to establish causality and investigate biological mechanisms, we undertook experiments in mice, finding that offspring exposed to NNS *in utero* had increased adiposity compared to controls, consistent with our observation in the CHILD cohort.

Our results add to an emerging body evidence from rodent studies examining early-life NNS exposure. Collison et al. (23) showed that exposing mice to aspartame *in utero* and throughout life (55 mg/kg/day, 1.4-fold ADI) resulted in increased body weight, visceral fat deposition, and fasting glucose levels, while von Poser Toigo et al. (25) found that male rat offspring exposed to high levels of aspartame (343 mg/kg/day, 8.6-fold ADI) during gestation had increased weight gain. In contrast, Olivier-Van Stichelen et al. found that maternal NNS throughout pregnancy and lactation either had no impact (for sucralose combined with acesulfame-K at levels approximating the ADI) or reduced offspring body weight (for higher doses of ∼2-fold ADI) (24), although adiposity was not measured and the offspring were not followed beyond weaning. Here, we separately assessed physiologically-relevant doses of aspartame and sucralose consumption. We showed that exposures approximating the human ADI of these NNSs during pregnancy and lactation increased body weight in male offspring, primarily due to an increase in their adiposity. Interestingly, and similar to sex-specific findings by Collison et al. (23), female offspring did not experience these effects. These findings are also consistent with sex differences observed in the CHILD infants at 1 year of age (14), although not replicated in our current analysis at 3 years of age. Further research is needed to understand the potentially sex-specific effects of NNS during critical periods of development.

We also uniquely evaluated the impact of maternal NNS intake on glucose and insulin tolerance in the offspring. Previously, Collison et al. found that exposure to 55 mg/kg/day of aspartame throughout gestation and postnatal life increased fasting blood glucose levels in both male and female offspring and decreased insulin sensitivity in male offspring only (23). While we did not detect differences in fasting blood glucose in NNS-exposed offspring, we did observe greater insulin resistance in the male offspring, which was consistent with their obesity. Since insulin resistance typically precedes the development of glucose intolerance and hyperglycemia, it is possible that these phenotypes could develop with advanced age or the addition of a high calorie diet.

Finally, we used a cell culture model of adipocyte differentiation to further explore the mechanisms of NNS-induced adiposity observed in the CHILD cohort and mouse offspring. Previously it was reported that saccharin and aspartame affected adipocyte differentiation and lipid metabolism (27-29), but these studies used extremely high millimolar dosages. Since we observed the greatest effects of sucralose on male mouse offspring, we treated male 3T3-L1 pre-adipocyte cells with 200nM sucralose at different stages of the differentiation process. We found that sucralose exposure very early in the differentiation program had the greatest effect on increasing lipid accumulation within the cells. In addition, this treatment increased the expression of several transcription factors that convert pre-adipocytes into adipocytes and have key roles in the regulation of lipid and glucose metabolism by adipocytes (31). These include PPAR-γ and C/EBP-α, as well their downstream target genes *Adipoq, Fabp4* and *Plin2*. Moreover, sucralose stimulated the expression of several genes involved in lipid metabolism, including *Fasn, Gpam, Lipe* and *Atgl*. Importantly, we confirmed that these changes in gene expression were also present in adipose tissues isolated from male offspring exposed to sucralose *in utero*. Together, these findings suggest that sucralose can directly induce a pro-adipogenic gene expression program at doses that approximate human consumption.

The major strength of this study is our translational approach. We used data from a large, longitudinal national birth cohort that collected objective measures of body composition and accounted for many possible confounders. We performed complementary mechanistic studies in mice and cultured adipocytes, and assessed two different NNS at physiologically-relevant doses. Limitations of the CHILD cohort study include the limited assessment of NNS in beverages, without details on the type of NNS, or NNS in foods, which are an increasingly common source of NNS exposure. As in all observational studies, residual confounding is also possible, although we accounted for key factors including maternal BMI, diabetes and diet quality. To overcome these limitations, we used experimental models to address causality and examine mechanisms. A limitation of our mouse study is that we did not separate the effects of NNS during pregnancy and lactation. A limitation of our adipocyte differentiation study is that although we used a dose that is relevant to human consumption, sucralose is not fully absorbed from the gut (32); therefore, the dose we applied to our cell culture system might be higher than what is achieved *in vivo*. However, our *in vitro* results were confirmed in mouse adipose tissue, demonstrating the compatibility of these model systems. Overall, our findings from the CHILD cohort and the experimental model systems are complementary and provide new insights into the biological impact of prenatal NNS exposure.

Further research is needed to confirm and characterize the potentially sex-specific biological mechanisms by which prenatal NNS exposure influences postnatal weight gain and adiposity. In addition to stimulating adipocyte differentiation, NNS may alter the maternal microbiome (33, 34), which is transmitted to the offspring during birth and postnatal interactions (35, 36), and contributes to host metabolism and weight gain (37-39). Future studies should also assess other types and sources of NNS, such as plant-derived NNS and NNS in foods. Finally, it will be important to study and model the maternal conditions that motivate NNS use, notably obesity and gestational diabetes, to clearly establish and disentangle their independent effects on offspring development. This research will be important for establishing the long-term safety of prenatal NNS exposure, and informing recommendations for pregnant women.

In summary, our translational research provides new evidence that exposure to NNS *in utero* stimulates postnatal weight gain, insulin resistance, and adiposity. Associations observed in the CHILD cohort were investigated in experimental model systems, revealing a previously unknown mechanism involving altered expression of pro-adipogenic (e.g. *Cebpa*) and lipid metabolism genes (e.g. *Gpam, Fasn*). Collectively, these results suggest that maternal NNS consumption is a modifiable obesogenic exposure that may be contributing to the global obesity epidemic, and call for further research on the long-term metabolic effects of NNS exposure in early life.

## Materials and Methods

### CHILD birth cohort

We accessed data from the CHILD cohort, a national population-based pregnancy cohort of 3455 families across four sites in Canada, enrolled between 2008-12 (40). For the current study, we included 2298 mother-infant dyads with complete data on maternal NNS consumption and child BMI at 3 years of age. This study was approved by the Human Research Ethics Boards at the Hospital for Sick Children, McMaster University and the Universities of Manitoba, Alberta, and British Columbia.

### Maternal dietary assessment

Maternal sweetened beverage consumption during pregnancy was documented in the CHILD study using a food frequency questionnaire (FFQ) (41, 42) as described previously (14). NNS beverages included “diet soft drinks or pop” (1 serving = 12 oz or 1 can) and “artificial sweetener added to tea or coffee” (1 serving = 1 packet). Sugar-sweetened beverage (SSB) intake was similarly determined from consumption of “regular soft drinks or pop” and “sugar or honey added to tea or coffee” (1 serving = 1 teaspoon or 1 packet). Beverage intakes were classified according to the number of servings per week as: never or <1 per month, ≤1 per week, 2 – 6 per week, or ≥1 per day (43). Total energy intake (kcal/day) and the Healthy Eating Index (HEI-2010 score) (44) were derived from FFQ data using food composition tables from the University of Minnesota Nutrition Coding Center nutrient database (Nutrition Coordinating Center, Minneapolis, MN). Sweetened beverage consumption was assessed in children at 3 years of age. Since consumption was relatively infrequent, it was simply categorized as any vs. none.

### Child body composition

At 3 years of age, height, weight, and subscapular skin folds were measured by trained CHILD study staff following a standardized protocol. Age- and sex-specific z-scores were calculated against the 2006 World Health Organization reference.

### Covariates and confounders

Child sex, birth weight, gestational age, and maternal age were collected from hospital records. Maternal BMI was calculated from measured height and self-reported pre-pregnancy weight (14). Maternal smoking during pregnancy and education (as an indicator of socioeconomic status) was determined via questionnaire. Diabetes during pregnancy (gestational or otherwise) was determined from hospital records and self-report. Breastfeeding duration was reported by a standardized questionnaire. Child screen time was measured in hours per day as an indicator of physical inactivity. Fresh and frozen food consumption were ascertained via questionnaire as a basic indicator of child diet quality.

### Experimental mouse model

All procedures were approved by the Animal Welfare Committee of the University of Manitoba, which adheres to the principles for biomedical research involving animals developed by the Canadian Council on Animal Care and the Council for International Organizations of Medical Sciences. All mice were given ad libitum access to chow diet and water. Food and water consumption was monitored throughout the experiment. Male and female C57BL6J mice were obtained at 8 weeks of age from the University of Manitoba colony and were mated. Following mating, chow-fed dams were randomly assigned to drinking water (control), sucrose (45 g/L, ∼7.2 g/kg body weight/day anticipating 4 mL water intake and 25 g body weight), aspartame (0.2 g/L, ∼32 mg/kg body weight/day) or sucralose (0.04 g/L, ∼6.4 mg/kg body weight/day) throughout pregnancy and lactation. In a preliminary dose-finding study, dams received low (0.05g/L), medium (0.1g/L) and high (0.2g/L) levels of aspartame or alternatively low (0.01g/L), medium (0.02g/L) and high levels (0.04g/L) of sucralose consumption (**Table S6**). These NNS concentrations are relevant to human consumption as they translated to doses near or below the acceptable daily intake limits for humans (40 mg/kg for aspartame and 5 mg/kg for sucralose).. Dams were allowed to deliver naturally and at birth and when necessary, litters were reduced to eight pups (i.e.-4 males and 4 females) to avoid competition for food. Beginning at 3 weeks of age (the usual weaning age for mice), offspring were fed regular chow and tap water. Food and water intake and body weight were measured weekly for all offspring. At 12 weeks of age, offspring were anesthetized by intraperitoneal injection of a sodium pentobarbital overdose and blood was collected by cardiac puncture. Tissues were dissected, rinsed in PBS, weighed, and either fixed in 10% formalin or freeze clamped in liquid nitrogen and stored at −80°C for future analyses. For all analyses, data from cagemates were averaged and the litter was used as the unit of analysis (i.e.- n=6 litters per group were generated).

### Dual-Energy X-ray Absorptiometry (DEXA)

DEXA scans were performed on 11 week-old mouse offspring at CHRIM (Children’s Hospital Research Institute of Manitoba) facilities. A technician blinded to the offspring experimental groups took all the images once mice were anesthetized using isoflurane gas and immobilized.

### Adipose tissue histology and morphometry

Histopathological preparations and hematoxylin/eosin (HE) staining were performed by the University of Manitoba Core Platform for Histology according to standard procedures. For the analysis of adipocyte size and number, the internal diameters of 80 consecutive adipocytes from 2 randomly selected fields on each HE stained slide were measured under light microscopy using a digital micrometer under 20x magnification and averaged.

### Evaluation of glucose and insulin tolerance

A glucose tolerance test (GTT) and insulin tolerance test (ITT) were performed on 11 week-old offspring, as described previously (45). For a GTT, mice were fasted overnight and injected intraperitoneally with a 50% glucose solution (2g/kg). Blood glucose concentrations were determined using an ACCU-CHEK advantage glucose meter (Roche Diagnostics) using blood collected from the tail and after a glucose injection (at 15, 30, 45, 60, 90 and 120 min). For the insulin tolerance test, human recombinant insulin was used to prepare an insulin-saline solution that was injected intraperitoneally (1mU/kg) after a 4h fast and blood glucose from the tail was measured at baseline and after insulin injection (at 30, 60, 90 and 120 min).

### Cell culture

3T3-L1 pre-adipocyte cells were differentiated as described previously (46). Two days post confluency (Day 0), the cells were stimulated with 1 μM dexamethasone, 1 μg/ml insulin and 0.5mM methylisobutyl-xanthine in 10% Fetal Bovine Serum(FBS)/Dulbecco’s Modified Eagle’s Medium media (Sigma). Cells were fed with fresh media every two days, with insulin and FBS on Day 2 and FBS alone from Day 4 until Day 8. Throughout adipocyte differentiation, 200nM sucralose was added at different stages of cellular development (**Fig. 3A**) until day 8. At Day 8, the cells reached full differentiation and samples were collected for analysis. After fixing cells with 10% formaldehyde for 2h at room temperature, washed with 60% isopropanol, lipid accumulation was evaluated by oil red O staining for 1h at room temperature and washed twice with distilled water. An EVOS digital inverted microscope (AMG) was used to capture microscopic pictures of the plates. Approximately 12 pictures were taken for every 100mm plate. 7 pictures were taken around the outside of the plate and 5 were taken around the center of the plate.

### Analysis of mRNA expression

RNA was isolated from tissues and cells using a QIAshredder column and further purified using the RNeasy kit (Qiagen, Valencia CA). For qPCR analysis, cDNA was synthesized using the Protoscript kit (NEB, Ipswich, MA, USA). The QuantiTect SYBR Green PCR kit (Qiagen, Valencia CA, USA) was used to monitor amplification of cDNA on an CFX96 real-time PCR detection machine (Bio-rad, Hercules, CA). Expression of genes was assessed in duplicate using 2-ΔΔCT and data was normalized by the geometric averaging of multiple control genes (47), including eukaryotic initiation factor 2a (eIF2a) and cyclophilin A as the reference genes that were constant across all groups of offspring. Primer sequences were validated and are reported in **Table S7**.

### Statistical analysis

Data from mouse and *in vitro* experiments are presented as mean (+/−) SEM. Differences in measurements performed among four groups were analyzed using one-way ANOVA and a Bonferroni post-test using GraphPad Prism v7.0 and RStudio version 1.0.143. For the CHILD cohort analysis, the distribution of covariates across categories of beverage consumption was examined by univariate analysis (χ2 test or analysis of variance). Associations between sweetened beverage intake and child body composition were determined using multivariable regression. Models were mutually adjusted for maternal and child consumption of both beverage types and adjusted in a stepwise manner for pregnancy and early life covariates, three-year covariates, and maternal BMI and child sweetened beverage consumption. Results are presented as crude and adjusted β estimates and odds ratios with 95% confidence intervals. Stratified analyses were performed to assess potential sex differences. All tests were 2-sided, and statistical significance was set at P<0.05.

## Acknowledgements

We are grateful to all the families who took part in the CHILD study, and the whole CHILD team, which includes interviewers, nurses, computer and laboratory technicians, clerical workers, research scientists, volunteers, managers, and receptionists. We also acknowledge the excellent technical work of Mario Fonseca and Bo Xiang (University of Manitoba) and critical review by Shirin Moossavi (University of Manitoba).

## Author Contributions

MBA and VWD conceived of the study design, obtained funding for this research, and drafted the manuscript. MRS, PS, TJM, SET, PJM, and ABB obtained funding for and oversaw recruitment of the CHILD cohort and data collection. AA performed the statistical analysis of clinical data from the CHILD cohort under the supervision of MBA. RJS contributed nutritional expertise. MMT, AH, and KGC performed mouse and cell culture experiments under the supervision of VWD. All authors critically reviewed and approved the manuscript.

## Data and materials availability

Data are available upon request from the corresponding authors. The CHILD Study data access policy is available at https://childstudy.ca/for-researchers/data-access/.

## Supplementary Materials

**Supplemental Table S1.** Maternal and child characteristics according to maternal consumption of NNS-sweetened beverages during pregnancy in the CHILD birth cohort.

|  | Frequency of NNS-Sweetened Beverage Consumption<br>During Pregnancy |  |  |  | p* |
| --- | --- | --- | --- | --- | --- |
|  | < 1 per month<br>N=2125 (70.1%) | ≤ 1 per week<br>N=512 (16.9%) | 2 - 6 per week<br>N=237 (7.8%) | ≥ 1 per day<br>N=159 (5.2%) |  |
| **Mother during pregnancy** |  |  |  |  |  |
| Age, years | 32.3 ± 4.7 | 32.6 ± 4.9 | 32 ± 4.4 | 32.1 ± 4.9 | 0.35 |
| BMI, kg/m<sup>2</sup> | 24.2 ± 5.0 | 25.6 ± 5.7 | 26.7 ± 6.7 | 28.1 ± 6.6 | <0.001 |
| Energy intake, calories/day | 2015 ± 715 | 2005 ± 709 | 2070 ± 816 | 2108 ± 907 | 0.30 |
| Diet quality, HEI Score | 73.0 ± 8.7 | 73.5 ± 8.1 | 71.8 ± 9.4 | 71.2 ± 8.6 | 0.005 |
| Smoking | 7.7% | 8.9% | 14.4% | 17.3% | <0.001 |
| Diabetes | 4.9% | 7.4% | 9.3% | 15.3% | <0.001 |
| No Postsecondary Degree | 22.4% | 21.1% | 29.4% | 28.8% | 0.02 |
| **Child at birth / during infancy** |  |  |  |  |  |
| Infant gestational age, weeks | 39.2 ± 1.4 | 39.2 ± 1.4 | 39 ± 1.5 | 39 ± 1.3 | 0.09 |
| Infant birth weight, g | 3451 ± 484 | 3446 ± 479 | 3403 ± 543 | 3463 ± 437 | 0.54 |
| Breastfeeding duration <12 mo | 52.8% | 60.5% | 69.0% | 75.7% | <0.001 |
| **Child at 3 years** |  |  |  |  |  |
| Screen time exceeds 1h/day | 76.2% | 81.9% | 83.8% | 86.3% | 0.003 |
| Fruit/vegetables <5/day | 60.9% | 58.8% | 61.7% | 61.2% | 0.88 |
| Fresh food <11/wk | 36.2% | 42.5% | 43.8% | 48.0% | 0.01 |
| Frozen food >3/wk | 55.6% | 62.4% | 65.6% | 71.4% | <0.001 |
| Boxed/processed food >3/wk | 42.0% | 44.1% | 48.8% | 57.1% | 0.01 |

Values are mean ± standard deviation for continuous variables, or percentages for binary variables.
\*Test for difference across NNS consumption categories, by ANOVA for continuous variables or chi-squared for binary variables.
HEI, healthy eating index 2010 (maximum 100); NNS, non-nutritive sweeteners.

**Supplemental Table S2.** Maternal sweetened beverage consumption during pregnancy and child BMI z-score at 3 years of age in the CHILD cohort

| Maternal Beverage Consumption | N | Child BMI z-score Mean ± SD | Crude $\beta$ (95%CI) | Adjusted for Maternal BMI | Model 1 Adjusted for Pregnancy/Early Life Covariates* | Model 2 Model 1 + 3y Covariates** | Model 3 Model 2 + child beverage intake and maternal BMI |
| --- | --- | --- | --- | --- | --- | --- | --- |
|  |  | N=2298 | N=2298 | N=2230 | N=2254 | N=1857 | N=1797 |
| **NNS Beverages** |  |  |  |  |  |  |  |
| < 1 per month | 2125 | 0.53 ± 0.96 | 0.00 (0.00, 0.00) | 0.00 (reference) | 0.00 (reference) | 0.00 (reference) | 0.00 (reference) |
| ≤ 1 per week | 512 | 0.60 ± 0.94 | 0.07 (-0.04, 0.17) | -0.01 (-0.12, 0.10) | 0.04 (-0.07, 0.15) | 0.07 (-0.05, 0.18) | 0.01 (-0.11, 0.13) |
| 2 - 6 per week | 237 | 0.64 ± 0.98 | 0.11 (-0.04, 0.26) | 0.04 (-0.11, 0.19) | 0.06 (-0.09, 0.21) | 0.07 (-0.11, 0.24) | 0.01 (-0.17, 0.19) |
| ≥ 1 per day | 159 | 0.88 ± 0.97 | **0.37 (0.19, 0.55)** | **0.23 (0.05, 0.42)** | **0.30 (0.12, 0.48)** | **0.26 (0.05, 0.48)** | 0.17 (-0.05, 0.39) |
| **SS Beverages** |  |  |  |  |  |  |  |
| < 1 per month | 671 | 0.49 ± 0.93 | 0.00 (reference) | 0.00 (reference) | 0.00 (reference) | 0.00 (reference) | 0.00 (reference) |
| ≤ 1 per week | 785 | 0.55 ± 0.93 | 0.08 (-0.04, 0.19) | 0.07 (-0.04, 0.18) | 0.06 (-0.06, 0.17) | 0.04 (-0.08, 0.17) | 0.05 (-0.08, 0.18) |
| 2 - 6 per week | 836 | 0.57 ± 0.96 | 0.09 (-0.02, 0.20) | 0.06 (-0.05, 0.18) | 0.06 (-0.05, 0.18) | 0.03 (-0.10, 0.16) | 0.02 (-0.11, 0.15) |
| ≥ 1 per day | 741 | 0.65 ± 1.00 | **0.17 (0.05, 0.28)** | **0.13 (0.02, 0.25)** | 0.12 (-0.002, 0.24) | 0.10 (-0.03, 0.24) | 0.10 (-0.03, 0.24) |

NNS, non-nutritive sweetener; SS, sugar-sweetened. BMI, body mass index; $\beta$ , beta estimate; CI, confidence interval; SD, standard deviation.
\*Pregnancy/ early life covariates include maternal total energy intake and Healthy Eating Index score, maternal postsecondary education, maternal smoking and diabetes in pregnancy, breastfeeding duration, and child sex. \*\*3y covariates include child screen time, fresh and frozen food intake, and soda consumption.
All models are mutually adjusted for both beverage types.

**Supplemental Table S3.** Maternal sweetened beverage consumption during pregnancy and child subscapular skinfold z-score at 3 years of age in the CHILD cohort

| Maternal Beverage Consumption | N | Child skinfold z-score Mean ± SD | Crude $\beta$ (95%CI) | Adjusted for Maternal BMI | Model 1 Adjusted for Pregnancy/Early Life Covariates* | Model 2 Model 1 + 3y Covariates** | Model 3 Model 2 + child beverage intake and maternal BMI |
| --- | --- | --- | --- | --- | --- | --- | --- |
|  |  | N=2003 | N=2003 | N=1946 | N=1967 | N=1632 | N=1580 |
| **NNS Beverages** |  |  |  |  |  |  |  |
| < 1 per month | 2125 | -0.01 ± 1.22 | 0.00 (reference) | 0.00 (reference) | 0.00 (reference) | 0.00 (reference) | 0.00 (reference) |
| ≤ 1 per week | 512 | 0.01 ± 1.14 | 0.03 (-0.12, 0.17) | -0.03 (-0.18, 0.11) | 0.02 (-0.13, 0.17) | 0.03 (-0.14, 0.19) | -0.01 (-0.18, 0.15) |
| 2 - 6 per week | 237 | 0.19 ± 1.32 | 0.21 (-0.01, 0.42) | 0.15 (-0.07, 0.36) | 0.19 (-0.03, 0.40) | 0.21 (-0.04, 0.45) | 0.18 (-0.07, 0.44) |
| ≥ 1 per day | 159 | 0.33 ± 1.32 | **0.35 (0.11, 0.60)** | 0.23 (-0.01, 0.48) | **0.29 (0.04, 0.54)** | 0.24 (-0.04, 0.53) | 0.17 (-0.13, 0.46) |
| **SS Beverages** |  |  |  |  |  |  |  |
| < 1 per month | 671 | -0.06 ± 1.24 | 0.00 (reference) | 0.00 (reference) | 0.00 (reference) | 0.00 (reference) | 0.00 (reference) |
| ≤ 1 per week | 785 | 0.09 ± 1.19 | 0.17 (0.02, 0.32) | 0.18 (0.02, 0.33) | 0.14 (-0.01, 0.30) | 0.15 (-0.02, 0.33) | 0.19 (0.01, 0.36) |
| 2 - 6 per week | 836 | -0.03 ± 1.22 | 0.05 (-0.10, 0.20) | 0.03 (-0.13, 0.18) | 0.02 (-0.14, 0.18) | -0.01 (-0.18, 0.16) | 0.01 (-0.17, 0.18) |
| ≥ 1 per day | 741 | 0.08 ± 1.25 | 0.15 (-0.01, 0.31) | 0.13 (-0.03, 0.29) | 0.08 (-0.09, 0.24) | 0.07 (-0.12, 0.25) | 0.11 (-0.08, 0.30) |

NNS, non-nutritive sweetener; SS, sugar-sweetened. BMI, body mass index; $\beta$ , beta estimate; CI, confidence interval; SD, standard deviation.
\*Pregnancy/ early life covariates include maternal total energy intake and Healthy Eating Index score, maternal postsecondary education, maternal smoking and diabetes in pregnancy, breastfeeding duration, and child sex. \*\*3y covariates include child screen time, fresh and frozen food intake, and soda consumption.
All models are mutually adjusted for both beverage types.

**Supplemental Table S4.** Average daily energy intake by mouse offspring at 11 weeks of age.

|  | Control | Sucrose | Aspartame | Sucralose |
| --- | --- | --- | --- | --- |
| **Males** |  |  |  |  |
| Food consumption (g) | 3.17 ± 0.07 | 3.59 ± 0.07 | 3.64 ± 0.09 | 3.52 ± 0.04 |
| Energy intake (kcal/day) | 56.4 ± 1.2 | 61.0 ± 1.2 | 61.8 ± 1.5 | 59.9 ± 0.5 |
| Energy intake (kcal/day/kg) | 2160 ± 42 | 2146 ± 26 | 2200 ± 41 | 2165 ± 31 |
| **Females** |  |  |  |  |
| Food consumption (g) | 3.30 ± 0.12 | 3.42 ± 0.03 | 3.22 ± 0.05 | 3.41 ± 0.24 |
| Energy intake (kcal/day) | 56.0 ± 2.1 | 58.2 ± 0.3 | 54.8 ± 0.9 | 57.8 ± 4.1 |
| Energy intake (kcal/day/kg) | 2496 ± 55 | 2543 ± 18 | 2546 ± 21 | 2744 ± 177 |

n=6 litters (4 male and 4 female offspring) per group. Values are mean ± SEM.
Comparisons by One-way ANOVA and Bonferroni post-test. \*p<0.05 vs Control (no significant differences).

**Supplemental Table S5.** Dose effects of maternal sucralose intake during pregnancy on mouse offspring body and tissue weights at 11 weeks of age

|  | Control:<br>No sucralose | Low<br>0.01 g/L | Medium<br>0.02 g/L | High<br>0.04 g/L |
| --- | --- | --- | --- | --- |
| **Males** |  |  |  |  |
| Body Weight (g) | 24.9 ± 0.5 | 26.0 ± 0.9 | 25.9 ± 0.4 | 27.2 ± 0.4* |
| gWAT (g) | 0.34 ± 0.01 | 0.34 ± 0.05 | 0.37 ± 0.05 | 0.46 ± 0.04* |
| pWAT (g) | 0.13 ± 0.01 | 0.09 ± 0.01 | 0.14 ± 0.02 | 0.21 ± 0.02* |
| Liver (g) | 1.17 ± 0.04 | 1.27 ± 0.07 | 1.21 ± 0.03 | 1.31 ± 0.08 |
| Heart (g) | 0.14 ± 0.01 | 0.14 ± 0.01 | 0.14 ± 0.01 | 0.15 ± 0.01 |
| Kidney (g) | 0.17 ± 0.01 | 0.17 ± 0.01 | 0.17 ± 0.01 | 0.17 ± 0.01 |
| Spleen (g) | 0.09 ± 0.01 | 0.07 ± 0.01 | 0.07 ± 0.01 | 0.09 ± 0.01 |
| **Females** |  |  |  |  |
| Body Weight (g) | 21.2 ± 0.6 | 20.7 ± 0.6 | 20.7 ± 0.4 | 21.1 ± 0.5 |
| gWAT (g) | 0.24 ± 0.03 | 0.22 ± 0.02 | 0.24 ± 0.03 | 0.22 ± 0.03 |
| pWAT (g) | 0.16 ± 0.02 | 0.12 ± 0.02 | 0.14 ± 0.02 | 0.16 ± 0.02 |
| Liver (g) | 0.89 ± 0.05 | 1.04 ± 0.03 | 0.99 ± 0.05 | 1.02 ± 0.05 |
| Heart (g) | 0.12 ± 0.01 | 0.11 ± 0.01 | 0.11 ± 0.01 | 0.12 ± 0.01 |
| Kidney (g) | 0.13 ± 0.01 | 0.13 ± 0.01 | 0.12 ± 0.01 | 0.14 ± 0.01 |
| Spleen (g) | 0.09 ± 0.01 | 0.09 ± 0.01 | 0.08 ± 0.01 | 0.09 ± 0.01 |

n=3 litters (4 male and 4 female offspring) per group. Values are mean ± SEM.
Comparisons by One-way ANOVA and Bonferroni post-test. \*p<0.05 vs Control

**Supplemental Table S6.** Tissue weights of mouse offspring at 11 weeks of age.

|  | Control | Sucrose | Aspartame | Sucralose |
| --- | --- | --- | --- | --- |
| **Males** |  |  |  |  |
| gWAT (g) | 338.3 ± 10.5 | 520.0 ± 61.4* | 524.8 ± 39.8* | 462.4 ± 35.7* |
| pWAT (g) | 127.4 ± 9.7 | 241.2 ± 44.7* | 235.8 ± 30.5* | 207.5 ± 24.4* |
| Liver (g) | 1.17 ± 0.04 | 1.26 ± 0.04 | 1.27 ± 0.03 | 1.31 ± 0.08 |
| Heart (g) | 0.14 ± 0.01 | 0.13 ± 0.01 | 0.15 ± 0.02 | 0.15 ± 0.01 |
| Kidney (g) | 0.17 ± 0.01 | 0.17 ± 0.01 | 0.17 ± 0.01 | 0.17 ± 0.01 |
| Spleen (g) | 0.09 ± 0.01 | 0.09 ± 0.01 | 0.08 ± 0.01 | 0.09 ± 0.01 |
| **Females** |  |  |  |  |
| gWAT (g) | 295.9 ± 38.2 | 415.3 ± 36.8* | 295.0 ± 18.1 | 268.9 ± 17.8 |
| pWAT (g) | 198.8 ± 41.8 | 330.0 ± 20.3* | 184.0 ± 15.6 | 199.1 ± 14.9 |
| Liver (g) | 0.89 ± 0.05 | 1.06 ± 0.04* | 0.99 ± 0.03 | 1.02 ± 0.05 |
| Heart (g) | 0.12 ± 0.01 | 0.11 ± 0.01 | 0.11 ± 0.01 | 0.12 ± 0.01 |
| Kidney (g) | 0.13 ± 0.01 | 0.13 ± 0.01 | 0.12 ± 0.01 | 0.14 ± 0.01 |
| Spleen (g) | 0.09 ± 0.01 | 0.09 ± 0.01 | 0.09 ± 0.01 | 0.09 ± 0.01 |

n=6 litters (4 male and 4 female offspring) per group. Values are mean ± SEM.
Comparisons by One-way ANOVA and Bonferroni post-test. \*p<0.05 vs Control.

**Supplemental Table S7.**
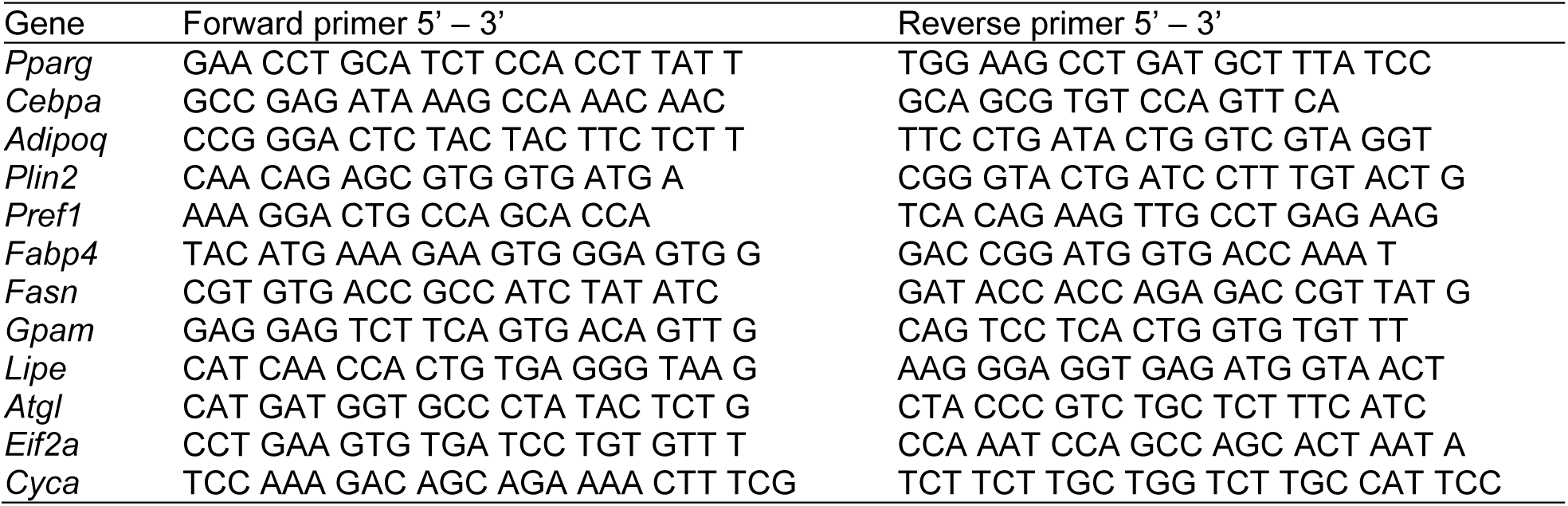
Primer sequences used for quantitative real-time PCR.

